# Haplotype phasing of *CYP2D6*: an allelic ratio method using Agena MassARRAY data

**DOI:** 10.1101/2023.02.27.530342

**Authors:** Megana Thamilselvan, Cheryl Mather, Yabing Wang, Katherine J. Aitchison

## Abstract

Pharmacogenomics aims to use the genetic information of an individual to personalize drug prescribing. There is evidence that pharmacogenomic testing before prescription may prevent adverse drug reactions, increase efficacy, and reduce cost of treatment. *CYP2D6* is a key pharmacogene of relevance to multiple therapeutic areas. Indeed, there are prescribing guidelines available for medications based on CYP2D6 enzyme activity as deduced from *CYP2D6* genetic data. The Agena MassARRAY system is a cost-effective method of detecting genetic variation that has been clinically applied to other genes. However, its clinical application to *CYP2D6* has to date been limited by weaknesses such as the inability to determine which haplotype was present in more than one copy for individuals with more than two copies of the *CYP2D6* gene. We report application of a new protocol for *CYP2D6* haplotype phasing of data generated from the Agena MassARRAY system. For samples with more than two copies of the *CYP2D6* gene for which the prior consensus data specified which one was present in more than one copy, our protocol was able to conduct *CYP2D6* haplotype phasing resulting in 100% concordance with the prior data. In addition, for three reference samples known to have more than two copies of *CYP2D6* but for which the exact number of *CYP2D6* genes was unknown, our protocol was able to resolve the number for two out of the three of these, and estimate the likely number for the third. In addition, we demonstrate that our method is applicable to *CYP2D6* hybrid tandem configurations.

## Introduction

Pharmacogenomics aims to use the genetic information of an individual to personalize drug prescribing.^1^ There is evidence that pharmacogenomic (PGx) testing before prescription may increase efficacy and reduce cost of treatment.^2-5^ Notably, a one-time genetic test can be cost effective in preventing adverse drug reactions.^6^ Pharmacogenes are those that affect the absorption, distribution, metabolism, and excretion of drugs, dietary substances, and toxins, as well as how these entities affect an organism.^1^ A study of about 500,000 participants in the UK Biobank looked at 14 different pharmacogenes and found that 99.5% of participants may show an atypical response to at least one drug; the average individual showing an atypical response to about 10 drugs.^7^ This implies that nearly everyone could benefit from PGx testing.

*CYP2D6* is a key pharmacogene of relevance not only for psychiatry,^8,9^ but also for other therapeutic areas.^10-12^ With over 150 different catalogued haplotypes, it is one of the best studied pharmacogenes.^13^ Moreover, as the majority of the variance in CYP2D6 enzyme activity has been shown to be genetically determined,^8^ and the functional result in terms of CYP2D6 enzyme activity of most *CYP2D6* haplotypes is known,^13^ it is the most useful gene in which to accurately characterize genetic variation in order to predict drug metabolism. While it is best known for its role in drug metabolism, it is widely expressed in multiple organs including the brain, and has physiological roles.^14^

*CYP2D6* is found adjacent to *CYP2D7* and *CYP2D8*, the latter two being pseudogenes.^15^ *CYP2D6* has 97% exonic sequence similarity with *CYP2D7*, and 92% with *CYP2D8*.^16^ The adjacent pseudogenes with high homology, along with intergenic repetitive sequences,^15,17^ predispose the region to the generation of a hypervariable locus.^18^ Indeed, with more than 150 different haplotypes,^13^ *CYP2D6* is one of the most variable human loci currently characterized. *CYP2D6* haplotypes include single nucleotide variants, insertions or deletions of short stretches of nucleotides (known as “indels”), and structural variants.^15^ Structural variants comprise duplications/multiplications, and deletions of the entire gene, as well as hybrid/fusion genes. Multiplications refer to at least 3 copies of the *CYP2D6* gene in tandem on one chromosome. Hybrid or fusion genes are those that are part *CYP2D6* and part *CYP2D7*.^15,18^ These include *CYP2D6-2D7* hybrids, in which the initial part of the gene is *CYP2D6* derived, followed by a *CYP2D7* derived region, or *vice versa* (*CYP2D7-2D6* hybrids).^15^ *CYP2D6-2D7* hybrids contain at least part of exon 9 from *CYP2D7*, and *CYP2D7-2D6* hybrids include at least exon 1 from *CYP2D7*.^15^ Hybrid genes can occur in more than one copy on one chromosome and also in tandem with another *CYP2D6* hybrid on one chromosome (known as hybrid tandems).

*CYP2D6* is located at chromosome 22q13.1.^19^ The combination of haplotypes on each of the two copies of chromosome 22 that an individual has is known as a diplotype. The overall resultant enzyme activity (or phenotype) has been categorized into four categories: poor, intermediate, normal, and ultrarapid metabolizers.^20^ Prescribing guidelines associated with phenotypes are available from the Clinical Pharmacogenetics Implementation Consortium (CPIC), the Dutch Pharmacogenetics Working Group (DPWG), the Canadian Pharmacogenomics Network for Drug Safety (CPNDS) and the French National Network (Réseau) of Pharmacogenetics (RNPGx).^21-23^

There are a variety of genetic technologies that aim to identify *CYP2D6* haplotypes. These include TaqMan Single Nucleotide Polymorphism (SNP) and Copy Number Variant (CNV) assays for *CYP2D6*, the Luminex xTAG CYP2D6 v3 kit, the Ion Ampliseq Pharmacogenomics Panel, PharmacoScan Solution, Agena Bioscience assays such as the Veridose Core Panel, and digital PCR.^24-30^ However, none of these at present claims to be able to conduct haplotype phasing for *CYP2D6*. There is one assay that was previously available and able to conduct haplotype phasing for a number of *CYP2D6XNs* (duplications/multiplications, specifically *\*1, *2, *4, *10, *17, *35*, and *\*41*), the AmpliChip CYP450 Test.^24^ Although this assay had this capability, it had weaknesses (as we and others have shown^24^), and was therefore withdrawn.

Haplotype phasing is required in the presence of *CYP2D6* duplications/multiplications, deletions, and hybrid tandems. For example, if a technology identifies that there is a *CYP2D6*1* haplotype and a *CYP2D6*4* haplotype and also more than one copy of one of these, it is necessary to know the phase (on which chromosome) the additional copy/copies lie. The *CYP2D6*1* haplotype is the wild-type (normal activity, assigned an enzyme activity score of 1), while *CYP2D6*4* is associated with zero enzyme activity; hence more than one copy of *CYP2D6*1* is associated with increased enzyme activity, while more than one copy of *CYP2D6*4* does not confer any additional enzyme activity. A *CYP2D6*1×2/*4* diplotype has an enzyme activity score of 2 (corresponding to a normal metabolizer), while a *CYP2D6*1/*4×2* has an activity score of 1 (corresponding to an intermediate metabolizer).

We herein present a method of haplotype phasing of *CYP2D6* based on calculation of the ratio of signals from one haplotype to another, for data generated using the Agena MassARRAY, using the Veridose Core Panel as an example. We show that this method works for *CYP2D6* duplications/multiplications and for hybrid tandems.

## Method

### Samples

DNA samples used were from the Genome-based therapeutic drugs for depression (GENDEP) study, comprising patients of European ancestry.^24^ The majority of these had prior data using the AmpliChip CYP450 Test (Roche Molecular Systems, Pleasanton, USA).^24^ In addition, positive controls from the Genetic Testing Reference Material Program (GeT-RM)^31^ were used: NA02016 (previously genotyped using the AmpliChip CYP450 Test as *CYP2D6*17/*2XN*), NA17221 (consensus *CYP2D6*1XN/*2*), and NA17439 (previously genotyped using the AmpliChip CYP450 Test as *CYP2D6*4XN/*41*).

### Data generation

The Agena MassARRAY system uses PCR amplification, followed by ionization of DNA and acceleration towards a detector,^32^ with the differential mass of ionized DNA molecules resulting in differential time to reach the detector and hence a mass spectrum. To date, the automated reports from the Veridose Core Panel and other pharmacogenetic panels have limitations including accuracy of estimation of number of duplicated/multiduplicated genes as well as haplotype phasing.^33,34^ It should be noted that the assay is not able to accurately identify haplotype copy numbers higher than three (=*X3* and above, denoted as 3N+ in the automated reports). The Veridose Core and Veridose CYP2D6 CNV Panels (Agena Bioscience, San Diego, U.S.A.) were run as per manufacturer recommendations, with a minor modification (adjustment of starting DNA template concentration, results best at 10 ng/μl). Samples were run on a MassARRAY Dx Analyzer, with data analysis being conducted using MassArrayTyper version 4.1, including PGx Report version 4.1.0.172. For CNV calls, the automated reports provide a functional CNV call (denoting the total number of functional *CYP2D6* haplotypes), as well as an overall CNV call (denoting the total number of *CYP2D6* copies). Using a method we developed,^35^ we conducted haplotype phasing by calculating the ratios of the peak heights of variant to reference alleles per SNP.

## Results

Data from the automated and allelic ratio adjusted genotype calls using our haplotype phasing method for the samples are presented in Table 1. For samples from the GENDEP set, exact copy number data were available in the prior consensus genotypes, and the adjusted genotype calls are 100% concordant with these. For the GeT-RM samples (for which the additional copies of the gene had previously been denoted just as “*XN*,” meaning that a duplication or multiplication was present, without identifying the number of *CYP2D6* gene copies), we were able to provide additional copy number information: specifically, that for NA17221 and NA17439, the *N* (or number of *CYP2D6* copies) is 2, i.e., two copies of the *CYP2D6*1* and *CYP2D6*4* haplotypes, respectively. For NA02016, as the ratio was 0.28, this likely approximates to 0.25, and hence 4 copies of the *CYP2D6*2* haplotype.

**Table 1.** *CYP2D6* automated and adjusted genotypes using the Veridose Core and Veridose CYP2D6 CNV Panels. Adjusted genotypes are derived from peak height variant to reference allelic ratios. The CNV number in the ‘Automated Call’ column is the functional CNV.

| Automated Call | Allelic ratios | Prior Consensus Genotype | Adjusted Genotype Call |
| --- | --- | --- | --- |
| 3N+ *2xN/*2xN | rs28371706: 0.28<br>rs16847: infinity<br>rs1135840: 24.62 | *2XN/*17 (NA02016) | *2X4/*17 or *2X3/*17 |
| 3N+ *1XN/*2XN | rs16947: 0.59<br>rs1135840: 0.66 | *2/*1XN (NA17221) | *2/*1X2 |
| 3N+ *4XN/*41XN | rs3892097: 2.2<br>rs28371725: 0.46 | *4XN/*41 (NA07439) | *4X2/*41 |
| 2N *1/*68+2 | rs16947: 0.59<br>rs1135840: 0.72 | *1X2/*2 | *1X2/*2 |
| 2N *2/*68+4 | rs16947: 2.09<br>rs1135840: 18.25<br>rs3892097: 0.61 | *2X2/*4 | *2X2/*4 |
| 3N+ *2XN/*68+*4XN | rs16947: 2.05<br>rs1135840: 15:05<br>rs3892097: 0.68 | *2X2/*4 | *2X2/*4 |
| 2N *1XN/*4XN | rs3892097: 2.24 | *4X2/*1 | *4X2/*1 |
| 3N+ *1XN/*2XN | rs16947: 0.63<br>rs1135840: 0.75 | *35/*1X2 | *2/*1X2 |
| 2N *1XN/*2XN | rs16947: 0.58<br>rs1135840: 0.77 | *2/*1X2 | *2/*1X2 |
| *68+*4/*4xN | rs1135840: 16.56<br>rs3892097: infinity<br>rs1065852: 11.75 | *4.013 +*4/*4 | *4.013 +*4/*4 |
| 2N *13 SNP FAIL | rs16947: 2.04<br>rs1135840: 1.65 | *13+*2/*1 | *13+*2/*1 |
| *2/*36xN+*10 | rs16947: 0.58<br>rs1135840: 15.95<br>rs1065852: 1.22 | (*36 + *10/*35) | Cannot conclusively determine based on SNP data only |

For the sample with the genotype *CYP2D6(*13+*2)/*1*, prior data aligned the *CYP2D6*13* to GQ162807 (or *CYP2D6*13A2*). As we have described, this haplotype is read as variant at rs16947 (2851C>T) and at rs1135840 (4181G>C) by other technologies.^24^ Therefore, two haplotypes on one chromosome (the *\*13* and the *\*2*) are variant at these positions, while the haplotype on the other chromosome is reference (*\*1*). Consistent with this, the ratio for rs16947 is 2.04, while the ratio for rs1135840 is 1.65. The mean calculated ratio for rs1135840 when the expected ratio is 2 (N=3) was 1.72 (95% CI [1.62, 1.81]). As the interpretation of the other hybrid tandems was less straightforward, we leave these to the Discussion.

## Discussion

We conclude using our protocol, it is possible to conduct haplotype phasing, and to determine which haplotype is present in more than one copy in data generated from the Agena MassARRAY system. Previous work has developed methods for haplotype phasing for SNP data generated from TaqMan and similar technologies,^36,37^ but to our knowledge this is the first report of a haplotype phasing method for *CYP2D6* data generated by the Agena MassARRAY system.

For evaluating the sample with the genotype *CYP2D6(*4*.*013+*4)/*4*, the defining SNP for *CYP2D6*4*, rs3892097, had a ratio of infinity, consistent the prior consensus genotype.^24^ The ratios for rs1135840 and rs1065852 are 16.56 and 11.75 respectively. There are some *CYP2D6*4* sub-haplotypes that have one or both of the SNPs, and some that have neither.^13^ The high ratio indicates that both *CYP2D6*4* genes present in this sample do have both of these SNPs; it is also consistent with the *CYP2D6*4*.*013* gene having the variant alleles or sequence variation in the region of these SNPs. For the sample with a consensus *CYP2D6* genotype of *(*36+*10)/*35*, the ratio for rs1135840 is 15.95. All three haplotypes (*CYP2D6*36, CYP2D6*10* and *CYP2D6*35)* are known to have the variant allele for this SNP, and the relatively high ratio is consistent with this.^13^ The ratio for rs16947 is 0.58, which can be approximated to 0.5. As *CYP2D6*35* has this SNP, but neither of the other two haplotypes do,^13^ the expected ratio is consistent with the consensus genotype. For rs1065852, the expected ratio is 2, as the *CYP2D6*10* and *CYP2D6*36* haplotypes currently catalogued by PharmVar both have the variant allele for this SNP, and *CYP2D6*35* does not.^13^ However, the calculated ratio was 1.22. The mean calculated ratio for rs1065852 in our samples where the expected ratio was 2 (N=3) was 1.29 (95% CI [1.21,1.37]), i.e., the calculated ratio tends to be lower than the expected ratio for this SNP. Therefore, the ratio of 1.22 is consistent with a *(*36+*10)/*35* genotype.

The discordant *CYP2D6*35* (*CYP2D6*1×2/*35* in the consensus genotype, *CYP2D6*2/*1×2* in our adjusted genotype) is owing to the fact that the *CYP2D6*35* haplotype is a variant of the *CYP2D6*2*, and the SNP discriminating *CYP2D6*35* from *CYP2D6*2* is not in the Veridose Core Panel. The function of both haplotypes is the same in the PharmVar database.^13^ For NA02016, four copies of *CYP2D6* have previously been described in specific ethnic groups, and the ethnicity of the sample is consistent with such reports.^38^ However, given the constraints of the genotyping technology at *CYP2D6* copy numbers of at least 3, it is possible that the ratio approximates to 0.33, and hence there are 3, not 4, copies of the *CYP2D6*2* haplotype. Another limitation of this allelic ratio technique for *CYP2D6* haplotype phasing is an inability to distinguish between various different possible genotypes where only one allele is present (e.g., C/C, CC/C, CC/- and C/-, or *\*1×2/*1* versus *\*1xN/*5*). Another limitation may be accuracy for certain SNP probes. For example, for rs201377835, which is the defining SNP for *CYP2D6*11*, and rs59421388, a defining SNP for *CYP2D6*29* and other haplotypes, we have seen non-zero values for the calculated allelic ratio for these SNPs where the consensus genotype does not include these haplotypes.

Nonetheless, given that the Agena MassARRAY system has demonstrated cost-effectiveness for clinical testing of other genes,^39,40^ and previously the inability to conduct haplotype phasing for *CYP2D6* represented a significant weakness that limited application of this technology to this gene, this paper represents an incremental contribution to pharmacogenomic testing for known variants in populations in which these have been identified. Should limitations such as the above be addressed, pre-emptive testing for *CYP2D6* prior to prescribing codeine or tramadol, for example,^41-43^ could prevent ineffective prescribing and adverse drug reactions.

## Acknowledgements

We thank Fang Yang at Alberta Precision Laboratories (APL) for running the Agena plates prepared by the Aitchison laboratory on the MassARRAY Dx system. We thank Xiuying Hu for participating in the Agena training, Keanna Wallace and Shui Jiang for general laboratory support, and Kinu Rill, Agena Bioscience field application scientist. The work described in this paper was funded in part by an Alberta Innovates Strategic Research Project (SRP51_PRIME - Pharmacogenomics for the Prevention of Adverse Drug Reactions in mental health; G2018000868 to KJA and Chad Bousman), a Canada Foundation for Innovation (CFI), John R. Evans Leaders Fund (JELF) grant (32147— Pharmacogenetic translational biomarker discovery, to KJA), an Alberta Centennial Addiction and Mental Health Research Chair (to KJA), and an Alberta Innovation and Advanced Education Small Equipment Grant (to KJA). GENDEP was funded by a European Commission Framework 6 grant, LSHB-CT-2003-503428. Roche Molecular Systems previously supplied the AmpliChip CYP450 Test arrays and some associated support, and we thank the GENDEP study participants for their contributions.

